# Speeding up Inference of Homologous Recombination in Bacteria

**DOI:** 10.1101/2020.05.10.087007

**Authors:** Felipe J Medina-Aguayo, Xavier Didelot, Richard G Everitt

**Affiliations:** Centro de Investigación en Matemáticas (CIMAT), Mexico; School of Life Sciences, University of Warwick, United Kingdom; Department of Statistics, University of Warwick, United Kingdom

## Abstract

Bacteria reproduce clonally but most species recombine frequently, so that the ancestral process is best captured using an ancestral recombination graph. This graph model is often too complex to be used in an inferential setup, but it can be approximated for example by the ClonalOrigin model. Inference in the ClonalOrigin model is performed via a Reversible-Jump Markov Chain Monte Carlo algorithm, which attempts to jointly explore: the recombination rate, the number of recombination events, the departure and arrival points on the clonal genealogy for each recombination event, and the range of genomic sites affected by each recombination event. However, the Reversible-Jump algorithm usually performs poorly due to the complexity of the target distribution since it needs to explore spaces of different dimensions. Recent developments in Bayesian computation methodology have provided ways to improve existing methods and code, but are not well-known outside the statistics community. We show how exploiting one of these new computational methods can lead to faster inference under the ClonalOrigin model.

## Introduction

Recombination is a critical process in evolution, particularly when analysing within-species variation. Bacteria reproduce clonally, but recombination exists in most species, where a donor cell contributes a small segment of its DNA to a recipient cell. This process is analogous to gene conversion in eukaryotes and is typically modelled using an Ancestral Recombination Graph (ARG) model (Hudson, 1990; Griffiths, 1996; Wiuf and Hein, 2000). The simulation of genomic data under this model is relatively easy (Didelot et al., 2009; Brown et al., 2016) but inference of the ancestral process given genomic data is much harder. This inference problem is important even when recombination is not directly of interest, because ignoring recombination leads to inaccurate reconstruction of the clonal part of the ancestry (Schierup and Hein, 2000; Hedge and Wilson, 2014).

The ClonalOrigin model (Didelot et al., 2010) can be regarded as a good approximation of the aforementioned ARG process, in which recombination events are modelled independently given the clonal genealogy, denoted throughout as 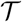. For completeness, we note the existence of related approximations to the ARG such as the sequential Markov coalescent for eukaryotes (McVean and Cardin, 2005; Marjoram and Wall, 2006) and the bacterial sequential Markov coalescent (De Maio and Wilson, 2017). There are also simpler approximations such as the ClonalFrame model (Didelot and Falush, 2007; Didelot and Wilson, 2015) in which the origin of each recombination event is not modelled.

The major challenge when implementing the ClonalOrigin model is to efficiently explore the joint posterior distribution of the recombination rate (*ρ*), the number of recombination events (*R*), their departures (*a*_1_,…,*a_R_*) and arrivals (*b*_1_,…,*b_R_*) on the clonal genealogy and the sites delimiting the start (*x*_1_,…,*x_R_*) and end (*y*_1_,…, *y_R_*) points of each recombination event on the genome. In order to explore such a complex distribution using Markov Chain Monte Carlo (MCMC), one must resort to the Reversible-Jump MCMC (RJMCMC) algorithm (Green, 1995) which in plain words aims at visiting spaces of different dimensions. This is the approach that was taken previously in both the original standalone implementation of the ClonalOrigin model (Didelot et al., 2010) and a recent reimplementation (Vaughan et al., 2017) within the BEAST2 framework (Bouckaert et al., 2019).

Unfortunately, as known by computational statisticians, the RJMCMC algorithm usually performs poorly due to the difficulty of proposing “good” trans-dimensional jumps. Because of this, the number of iterations (and consequently the running time) required by the algorithm for obtaining a reasonable approximation of the posterior distribution may be impractically large. Recent developments in Bayesian computation methodology provide ways of improving existing methods and code, but are not well-known outside the statistics community.

Such state-of-the-art methods belong to the framework of Sequential Monte Carlo (SMC) methods (see e.g. Doucet et al., 2001; Del Moral et al., 2006) where inference is performed using an appropriate sequence of intermediate target distributions leading to the desired one. Some of these methods have been successfully implemented to some extent when inferring coalescent trees as data arrives (see e.g. Dinh et al., 2018; Everitt et al., 2019). By performing inference in a sequential manner the complexity of the problem is reduced and the the desired target distribution may be explored more efficiently. The ideas presented here are based on the aforementioned SMC methodology that can lead to faster inference in the ClonalOrigin model; nonetheless, these ideas could be in principle applied to other evolutionary models.

## Methods

### Bayesian inference under the ClonalOrigin model

In this section we briefly describe the elements of the ClonalOrigin model, namely the parameters of interest, prior distributions on these parameters and the likelihood function for the data. For more details on assumptions and derivations of formulae please refer to cited references and the supplementary material.

- We use a coalescent tree to represent the clonal genealogy of *n* samples 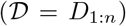 and denote such a tree by 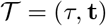, which is composed of a topology *τ* and a vector **t** = (*t*_2_,…,*t_n_*) of branch lengths. A Kingman’s coalescent prior (Kingman, 1982) is assumed for 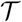.
- Let *R* denote the number of recombination events affecting the DNA sequences, which we assume a priori to be distributed according to a Poisson random variable with mean 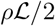, where *ρ* denotes the global recombination rate and 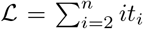 is the total branch length of the tree 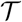, see e.g. Wiuf and Hein (1999) for more details. We also let 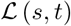 denote the sum of branch lengths from time *s* to time *t* on the tree.
- Each recombination event is formed by the following variables: departure and arrival points on the genealogy, and start and end sites on the genome. These four variables, when referring to the *i*-th recombination edge, are denoted by *a_i_, b_i_, x_i_* and *y_i_*, respectively.
- Both variables *a_i_* and *b_i_* are fully determined by a time and lineage on the tree (denoted respectively by *a_i_*[*t*] and *a_i_*[*l*], respectively), and the chosen priors are those from the construction of an ARG as a point process along the genomes (Wiuf and Hein, 1999).
- The priors for *x_i_* and *y_i_* are constructed assuming a uniform distribution on the sequence for *x_i_* and a geometric distribution of mean *δ* > 0 for the difference *y_i_* − *x_i_* | *x_i_*. When the sequence is made of *B* blocks comprising a total length of *L* the priors need to be modified accordingly as in Didelot et al. (2010).
- The entire set of variables describing the recombination events is denoted by

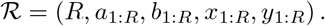
- Mutation events occur at rate *θ*/2 across the genealogy and on existing recombination edges; for simplicity we assume that all substitutions are equally likely, as in the evolutionary model JC69 (Jukes and Cantor, 1969).

Under the above the assumptions, the full prior for the set of parameters of interest is given by (see the supplementary material for the derivation)

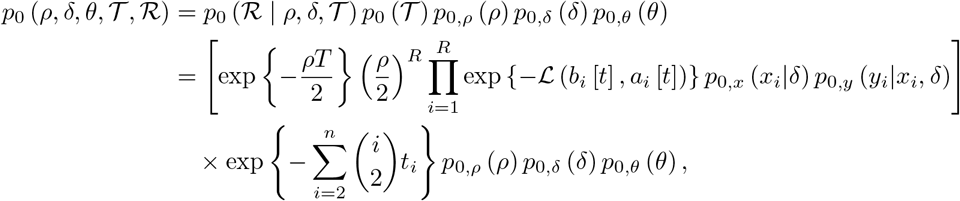

where *p*_0,*ρ*_,*p*_0,*δ*_,*p*_0,*θ*_ are arbitrary priors for the recombination rate, mean-tract length and mutation rate. For the likelihood computation, first recall that mutation events occur at rate *θ*/2 across the genealogy and on existing recombination edges; we then assume for simplicity that all substitutions are equally likely (Jukes and Cantor, 1969). The computation of the likelihood function 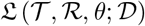, for any tree 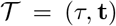, set of recombination events 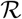, and mutation rate *θ*, is done using Felsenstein’s pruning algorithm (Felsenstein, 1973, 1981) (see also the supplementary material for more details).

The posterior on the full set of parameters is obtained through Bayes’ Theorem

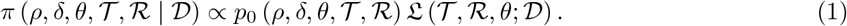

However, inferring the whole set of parameters represents a big challenge. One could instead fix one or more parameters and work with the resulting conditional distributions, for example finding a point estimate of 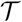 and then fix it to infer the rest of the parameters. We will consider this incomplete approach for some of the examples presented later where we aim to explore

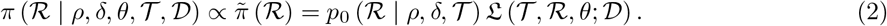

### The algorithm

Markov chain Monte Carlo (MCMC) is commonly the method of choice for exploring posterior distributions. If we had access to the the marginal posterior for the number of recombination events 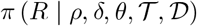, the inference of the remaining parameters could be carried out independently across different values of *R*. Since such a marginal is not available we need to infer *R* jointly with the rest of the parameters leading to a posterior that has no fixed dimension. The celebrated Reversible-Jump MCMC (RJMCMC) algorithm (Green, 1995) provides an elegant solution and corresponds to the generalisation of the Metropolis-Hastings (MH) algorithm that allows “jumps” across different dimensions. In our context, these jumps will correspond only to going up or down by one dimension, i.e. given *R* ≥ 1 recombination events they can go up to *R* + 1 or down to *R* − 1. In addition to these trans-dimensional moves, we still need to perform intra-dimensional moves that allow the full exploration of the desired posterior. Therefore, for fixed *R* we could also perform moves on 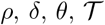 and 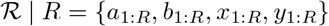.

Unfortunately, RJMCMC typically suffers from bad mixing in the sense that the resulting chain converges slowly to the desired posterior; this will in turn require a prohibitively large number of iterations for obtaining accurate answers. Due to recent developments in methodology (Karagiannis and Andrieu, 2013; Andrieu et al., 2018), the acceptance ratio for trans-dimensional moves within the RJMCMC can be understood as an importance sampling estimate of the ratio of two marginal densities, i.e. for an upwards move the ratio corresponds to 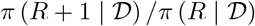, and using a single importance point (more details on this can be found in the supplementary material). One could then ask whether improving the aforementioned estimate could result in a chain with better mixing. The answer turns out to be positive but with some restrictions. Two straightforward approaches to achieve variance reduction of the estimator are annealed importance sampling (Neal, 2001) and simply using more than one importance point.

In plain words, the annealing procedure creates a smooth bridge between distributions of different dimension; so in a sense instead of attempting one big jump from *R* to *R* + 1 we attempt many, say *T* > 1, smaller jumps. Clearly, the downside of this approach is the extra cost of performing *T* steps that could result in a rejection in the MCMC algorithm. Whether annealing provides an advantage or not will entirely depend on the quality of the smaller jumps. In the ClonalOrigin context, when proposing to add a new recombination event that is in a bad region of the posterior, the idea is to use the smaller jumps to correct its position before deciding to accept or not.

On the other hand, considering *N* ≥ 2 importance points when jumping from *R* to *R* + 1 requires the creation of several proposed recombination events which are used for estimating the aforementioned ratio of marginals. However, if the upwards ends up being accepted one needs to select which of the proposed recombination events will be retained for the next iteration of the algorithm. To do this, a categorical distribution is used that selects which recombination event will survive according to its weight or contribution to the estimate of the acceptance ratio. Therefore, a recombination event that is more plausible than the rest will have a larger weight and, if the upwards move is accepted, it will have a greater chance of being retained ready for the next iteration.

Figure 1 illustrates both the annealing and the multiple importance points schemes when *T* = 5 and *N* = 4. The trees in black correspond to the current value of 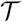 on which recombination events are appended. Colours correspond to different stages of the proposed recombination event when perturbed; the idea of annealing is that the proposed event in blue at *t* = 1 will be perturbed in such way that the the one in red at *t* = *T* has a better chance of being accepted. This process can be repeated *N* times, possibly in parallel, noting that perturbations could be drastic (as in *n* = *N* in the figure) or almost nonexistent (as in *n* = 3); it all depends on the quality of the the proposed event at *t* = 1 and the perturbation mechanism. Only one event from those at *t* = *T* is selected for the final accept-reject step in the MCMC, but events leading to a higher value in the posterior will have a better chance of being selected and possibly added later on, as discussed previously.

**Figure 1:**
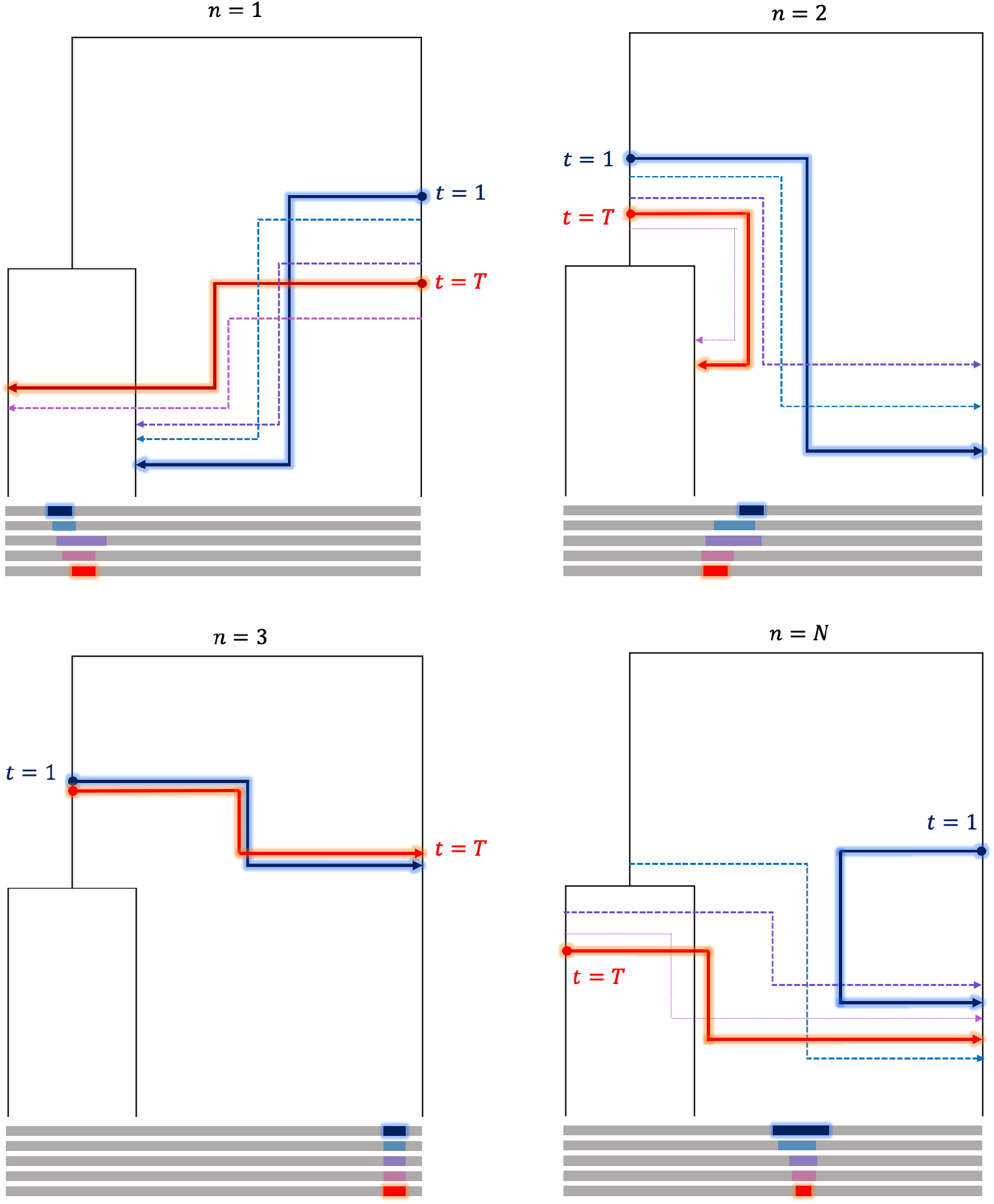
Illustration of annealing and multiple importance points for *T* = 5 and *N* = 4.

Algorithm 1 describes one step of the full process which we have termed Reversible-multiple-Jump MCMC (RmJMCMC) in line withB Andrieu et al. (2018), where it was firstly introduced and studied, and belonging to the wider class of MH with Asymmetric Acceptance Ratio (MHAAR) algorithms. Here we have adapted RmJMCMC to the ClonalOrigin context. We refer the reader to the appendix, where we describe the UAM and DAM algorithms used in the RmJMCMC. The implementation in `C++` of the RmJMCMC algorithm is freely available at: https://github.com/fmedina7/Clon0r_cpp

Notice that the upwards move agrees with the description in the previous paragraph; however, the downwards move appears to be more intricate. In words, we propose to delete a recombination event but in order to decide whether to accept or not we must test if this deletion is convenient. This is done by generating *N* − 1 recombination events and computing the resulting acceptance ratios as if we were trying an upwards move. Low values for the aforementioned ratios would imply that the proposed deletion is possibly a good decision, whereas if many ratios result in high values it might indicate that the deletion is a poor choice. The final decision rule results in a valid algorithm as shown in Andrieu et al. (2018), otherwise the algorithm would not be exact in the sense that it does not target the desired posterior distribution. Non-exact or noisy methods have been explored in the past (see e.g. Alquier et al., 2016), however we do not discuss them any further as the bias that is introduced is typically difficult to quantify.

One further observation is the obvious increased computational cost of RmJMCMC as opposed to performing only RJMCMC (equivalent to RmJMCMC when *T* = *N* = 1) or running multiple RJMCMC chains in parallel. Every iteration of RmJMCMC is at least *TN* times more expensive than one RJMCMC iteration. However, as opposed to running RJMCMC for longer or multiple independent RJMCMC chains, RmJMCMC has provable better convergence properties (Andrieu et al., 2018) that can reduce burn-in times and improve convergence towards a region of high posterior probability. This is discussed in more depth in the Results section where we look at some examples.

### Flavours of RmJMCMC

In order to implement Algorithm 1, we must specify the sampling auxiliary distributions 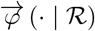 and 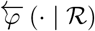 (which could both depend on other parameters, e.g. 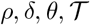). In the following section we present results using simple choices, the joint prior on (*a,b,x,y*) for the distribution 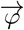 and a discrete uniform on the set {1,…, *R*} for 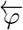 assuming there are *R* active recombination events. In mathematical terms, the associated densities are

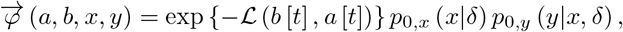

and 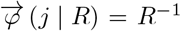 for *j* ∈ {1,…, *R*}. The previous choices greatly simplify the computation of the output ratios in Algorithms 2 and 3, which involve mainly ratios of likelihood functions.

#### Algorithm 1 Reversible-multiple-Jump MCMC (RmJMCMC)

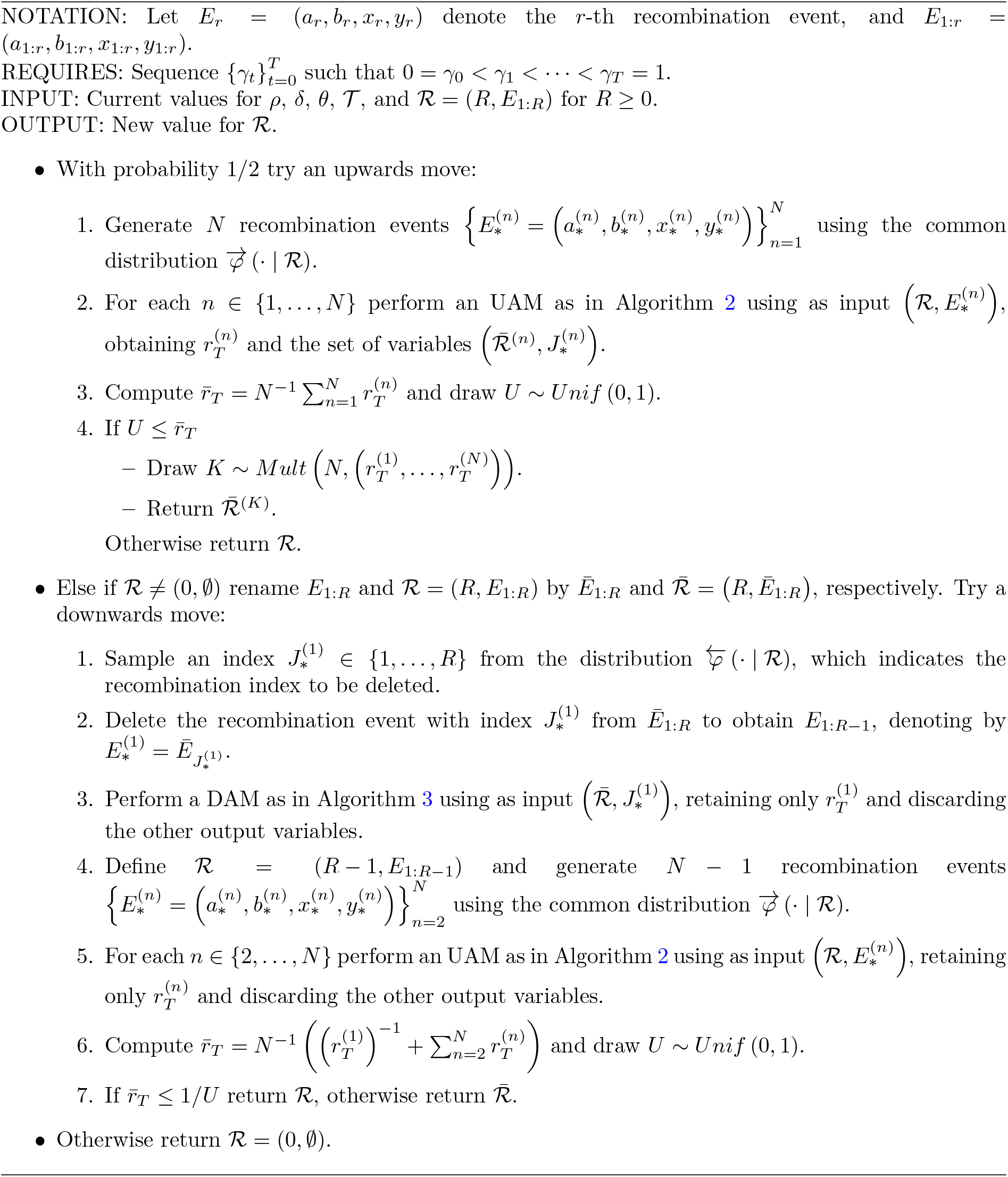

Additionally, we must define the way to perturb recombination events at every small step in the annealing process. This is done using an MCMC algorithm as explained in Algorithms 2 and 3; hence the perturbation is fully defined once we select a proposal distribution for 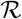 fixing the value of *R*, i.e. we need to perturb at least one recombination event within the existing *R* events. For the examples in the following section we choose to perturb (using the prior as proposal) only the newly created recombination event when going upwards, or the recombination event to be deleted when going downwards, for both cases this event is denoted by *E*\* in Algorithms 2 and 3. Doing this provides a very simple expression for the acceptance ratio in the MCMC steps within the annealing that involves only ratios of likelihood functions.

Finally, we must decide the number of annealing steps *T*, the sequence of real numbers 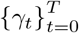 and the value of replicates *N*. In the examples that follow we use different values for *T* and *N* which involve different costs and running times. As mentioned earlier, the computational cost increased as *T* and *N* increases, however an appealing property of RmJMCMC is that loops involving *N* (either upwards or downwards) can be performed in parallel, this may lead to higher efficiency when computing running times. Due to this, the value of *N* is commonly determined by the number of cores available in the computer or server. Respecting the required sequence 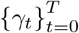 we simply choose a linear interpolation *γ_t_* = *t*/*T*.

We want to emphasise that the choice of 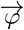 and 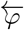 were made in accordance to the original ClonalOrigin implementation from Didelot et al. (2010); whereas the choices for the proposals within the MCMC and the elements of the sequence 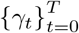 were made for convenience. However, the way RmJMCMC was formulated permits the use of more complex approaches that could provide better results in terms of efficiency. Some of these improvements are briefly described in the Discussion and are devoted to future work.

## Results

This section compares the RmJMCMC algorithm and the standard RJMCMC for the ClonalOrigin model. We start with some toy simulated experiments, moving later on to results using real, and previously studied, datasets. The code used for generating the forthcoming results is available at: https://github.com/fmedina7/Clon0r_cpp

### Application to simulated data

We present several simulated examples in order to explore the scalability and usability of RmJMCMC. We compare the settings when *N* = *T* =1 (corresponding to the RJMCMC algorithm), when *T* = 4 and *N* = 4, when *T* = 2 and *N* = 8, and when *T* =1 and *N* = 16. The efficiency between these configurations is compared using an estimation of the effective sample size (ESS) for a quantity of interest, as is commonly done in MCMC (Robert and Casella, 2004). In words, the ESS indicates how many independent samples were obtained from the total number of iterations for which the algorithm was run, and if the chain exhibits large correlation then the ESS will typically be much smaller than the number of iterations.

Throughout this section we consider a “typical setting” where data was simulated using a mean tract length of *δ* = 236, a mutation rate per site *θ_s_* = 0.03, and a recombination rate per site *ρ_s_* = 0.005. These three parameters remain fixed, and consequently the inference is carried only for the number of recombination events and their locations on the genome and on the tree, i.e. the target distribution is given by (2).

#### Example 1.

Typical setting: 50 sequences, length 50 kilobase pairs (Kb), mean tract length *δ* = 236, global mutation rate *θ* = 1500 (*θ_s_* = 0.03), global recombination rate *ρ* = 250 (*ρ_s_* = 0.005).

Figure 2 contains traceplots for the number of recombination events (top row) and for the value of the log-likelihood (middle and bottom rows). Observe that the schemes for which *N* > 1 have a similar performance when comparing against iteration number (plots on the left); this is not surprising since the annealing moves (those involving *T*) are perturbed using an MCMC algorithm with a proposal equal to the prior, and also the multiple instances (those involving *N*) are generated using the prior. However, the computational burden is different in each setting since for a fixed value of *NT* the algorithm could be parallelised using *N* cores but still requires *T* − 1 serial iterations of the annealing process. Hence, for this case, the setting where *T* =1 and *N* = 16 should perform best when taking into account the running time. The plots on the right incorporate this information and notice that all the schemes still seem to perform better than the standard RJMCMC chain (in blue) as they reach a region where the likelihood is high in a shorter amount of time, this can also be seen more clearly in the plots from the bottom row. It is worth noting that despite the RJMCMC chain being closer to the true number of recombination events, it is not a reliable indicator of whether the chain has converged since the value of the log-likelihood is still in a transient phase.

**Figure 2:**
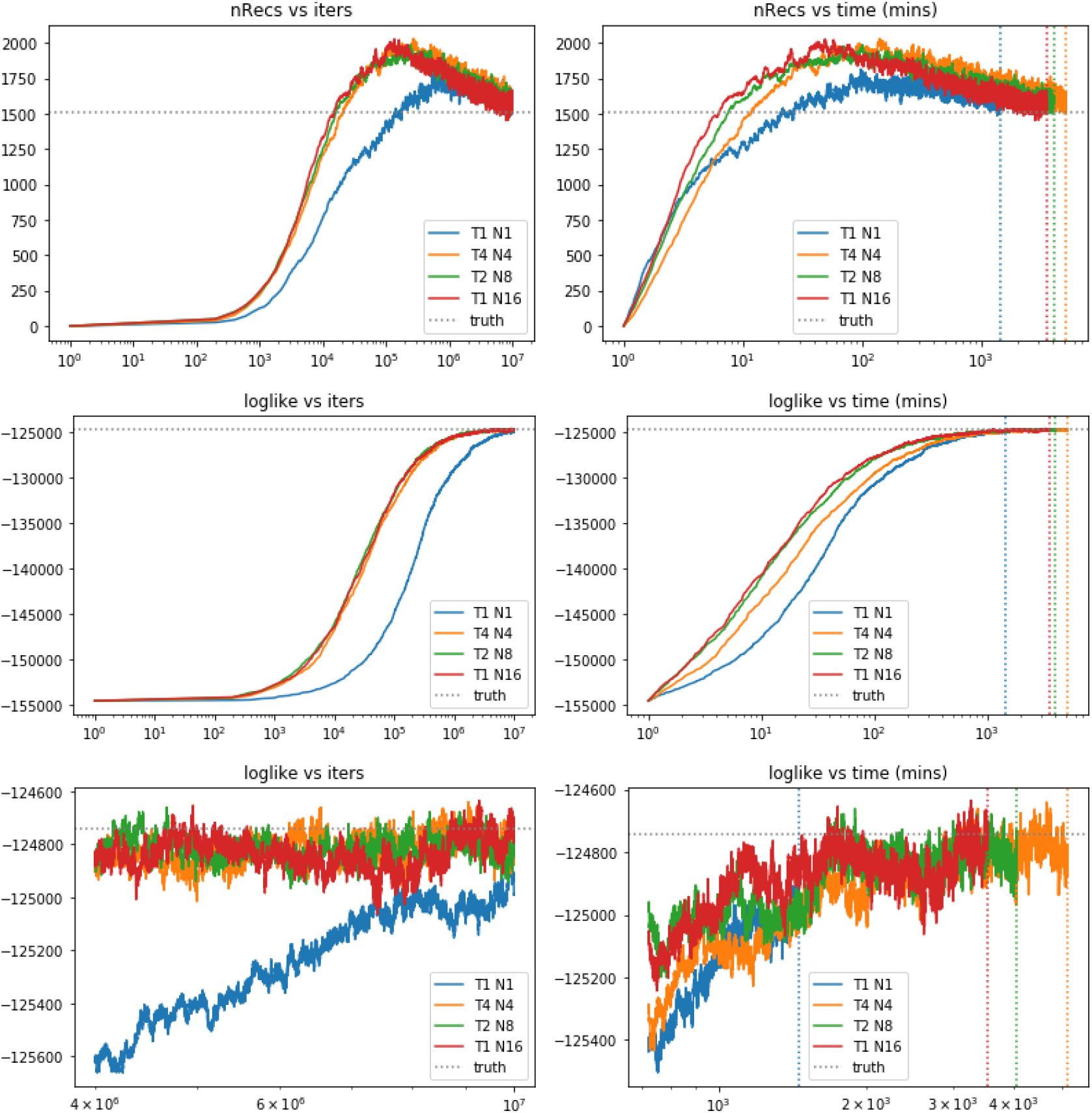
Trace plots for the total number of recombinations and for the log-likelihood for different combinations of *T* and *N* and using 10 million iterations. Plots on the left correspond to values vs iteration number, those on the right are vs running time. Grey dotted line corresponds to ground truth, coloured dotted lines indicate time when algorithm stopped. Bottom row: trace plots of the log-likelihood for the last 6 million iterations (left) and after 12 hours of running time (right).

Figure 3 compares the ESS fusing the values of the log-likelihood for the different schemes. Notice that when the running time is not taken into account (left plot) the three settings of RmJMCMC (those with NT = 16) have very similar ESS, as expected from the left plots of Figure 3. The right plot is in line with the conjecture that the setting when *T* = 1 and *N* = 16 is the most efficient, observing that all three still outperform the standard RJMCMC chain.

**Figure 3:**
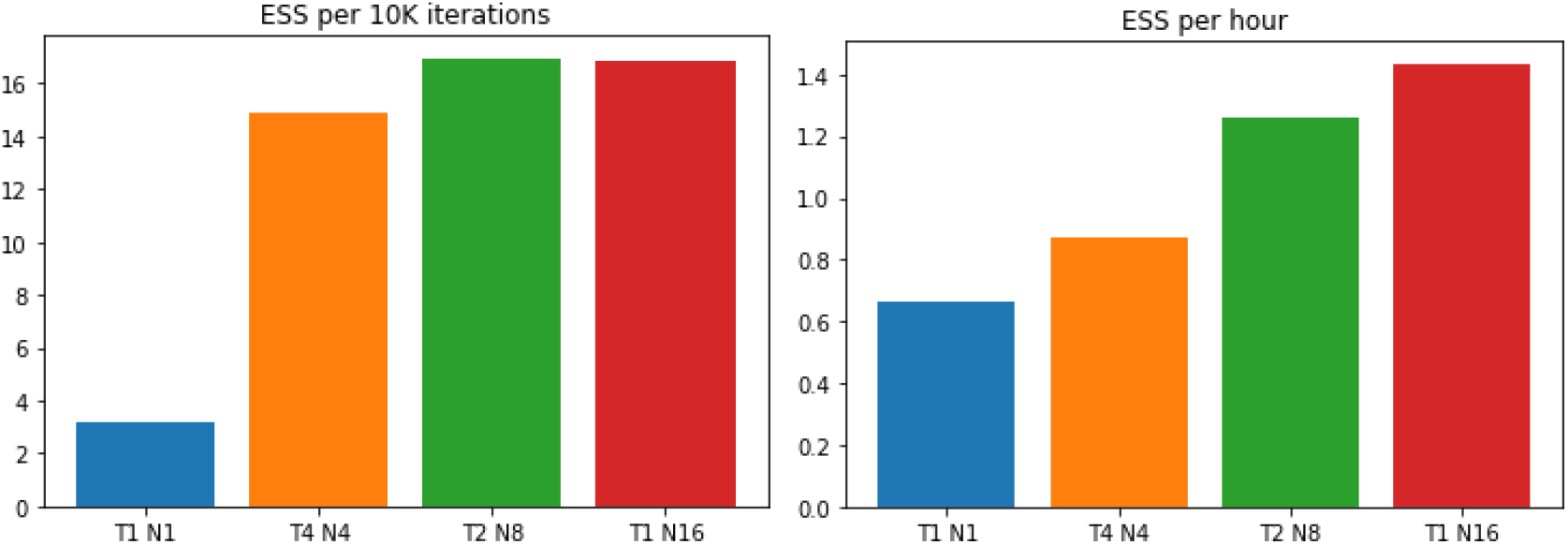
Effective sample sizes per 10 thousand iterations (left) and per hour of running time (right).

Figures 4 and 5 compare other quantities of interest. Figure 4 shows the recombination frequencies across the sequence of length 50K depicting the active number of recombinations affecting each site. Despite the apparent similarity of the results for the different schemes (top and middle rows), when looking at a smaller scale there are important deviances from the truth (bottom plots) that are direct consequence of a bad mixing from the chain. Settings with large values of *NT* have greater chances of better exploring the complicated state space in the ClonalOrigin model.

**Figure 4:**
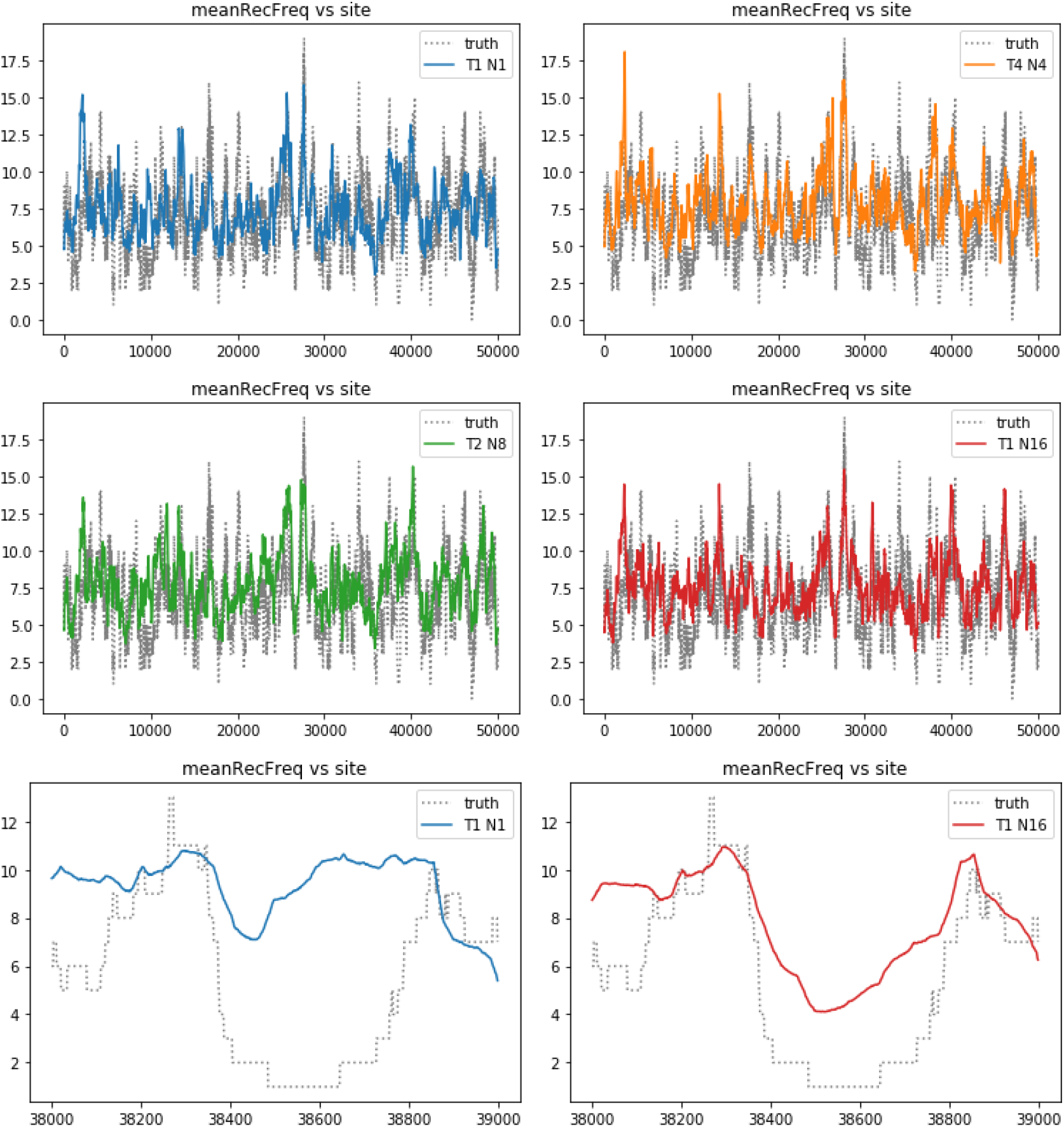
Average recombination frequency vs site number for different combinations of *T* and *N*. Grey dotted line corresponds to the true recombination frequency.

**Figure 5:**
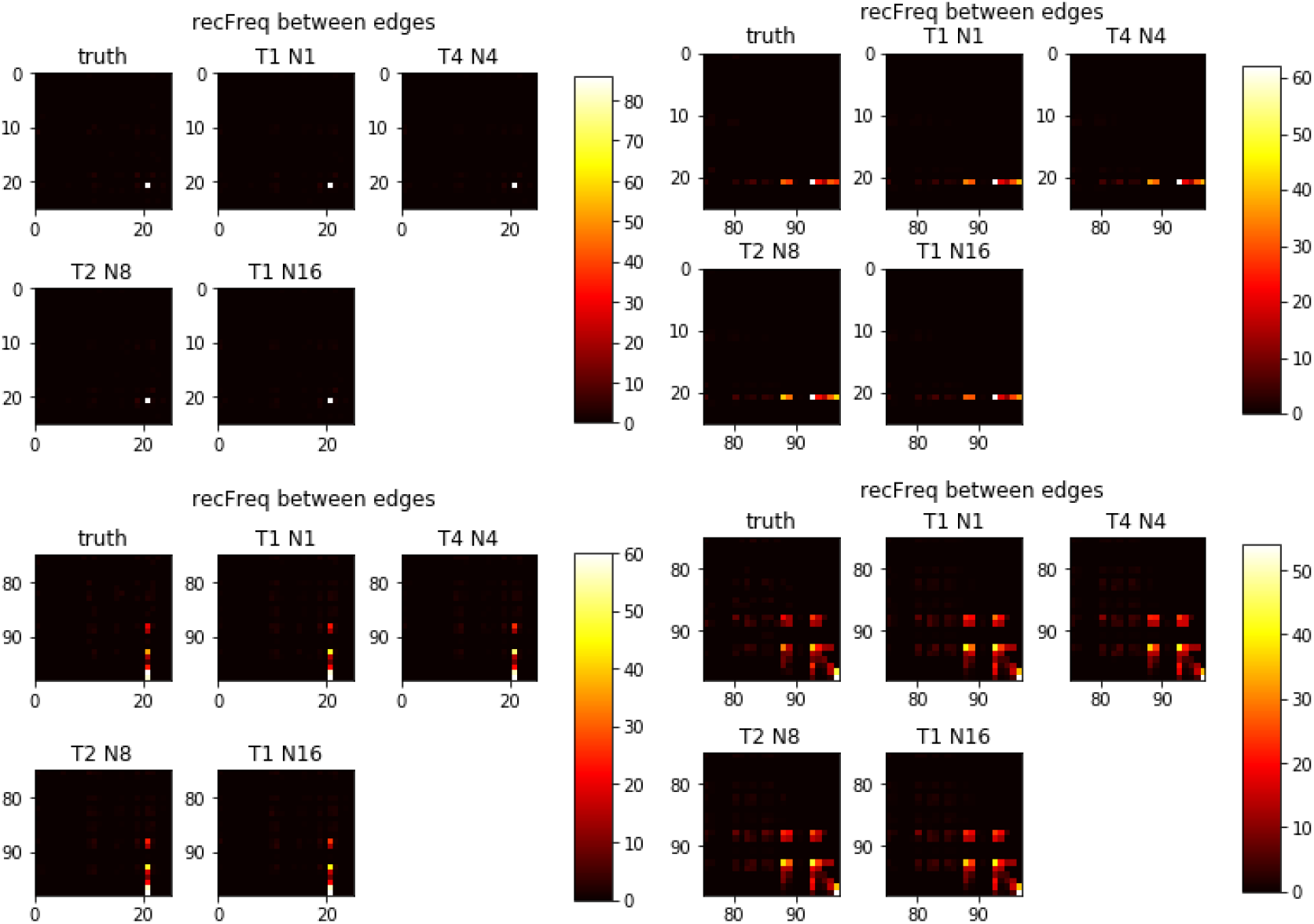
Average recombination frequency between edges, y-axis indicates the edge number where a recombination departed, x-axis denotes the edge number where a recombination landed. The plots correspond to the corners of the 98×99 matrices containing the recombination frequencies between edges.

Figure 5 also shows recombination frequencies, but between edges in the tree. Coloured squares indicate pair of edges on the tree joined by several recombination events, depending on the scale. In this case, the differences between the schemes and against the ground truth are negligible.

#### Example 2.

Typical setting: 50 sequences, length 10Kb, mean tract length *δ* = 236, global mutation rate *θ* = 300 (*θ_s_* = 0.03), global recombination rate *ρ* = 50 (*ρ_s_* = 0.005).

This example is similar to the previous one, except for the length of the sequence which is now 10Kb. The global mutation and recombination rates have been scaled appropriately leaving the per-site rates the same as before. Figure 6 contains some plots of interest, we observe once more an improvement of the ESS and faster convergence of the log-likelihood without considering running times. However, when time is taken into account (right plots) running RmJMCMC seems to be worth it only when *T* = 1 and *N* =16. Having shorter sequences makes the inference problem less expensive, so the schemes where *T* > 1 do not seem to add any benefit.

**Figure 6:**
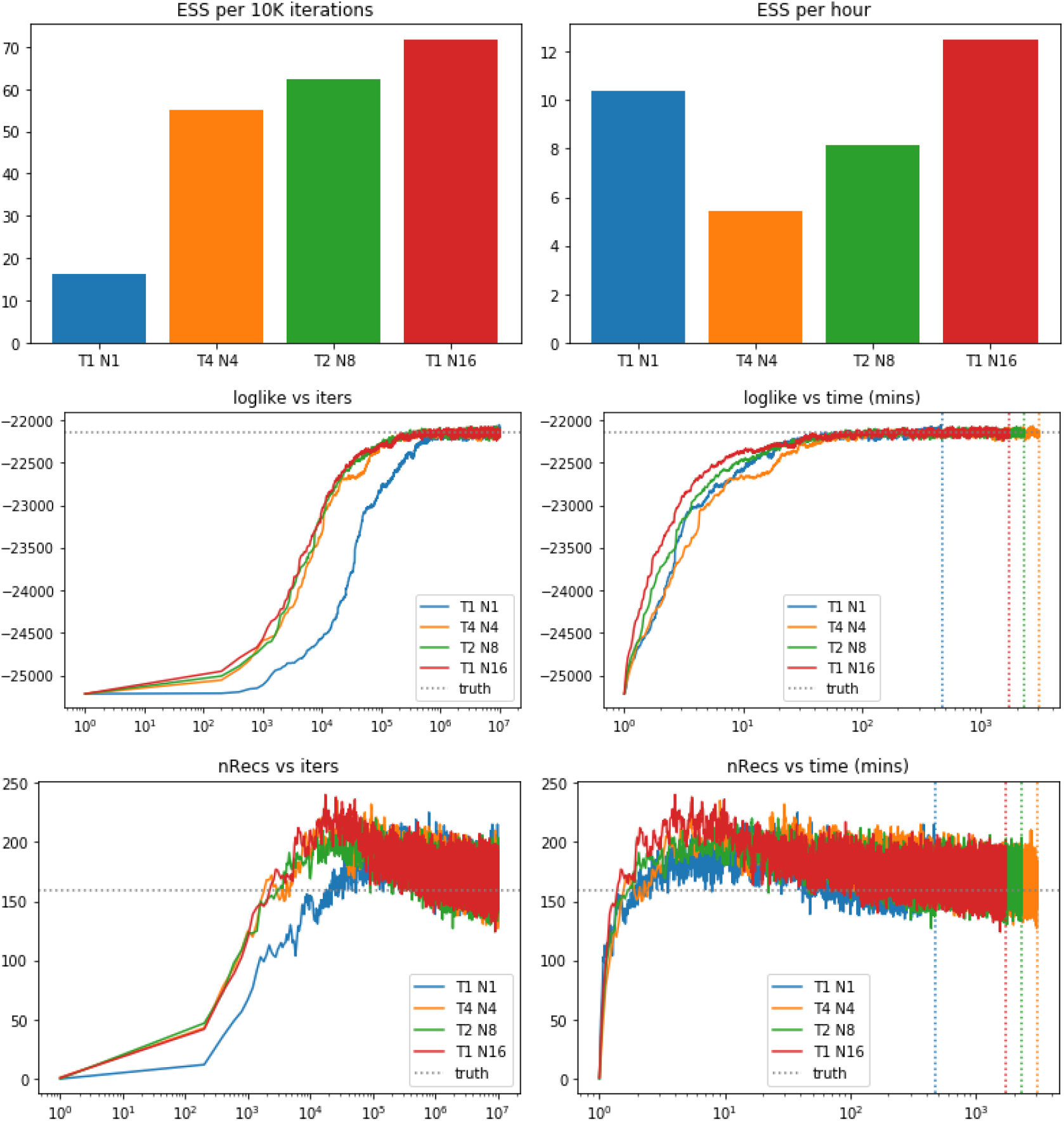
Top row: Effective sample sizes per 10 thousand iterations (left) and per hour of running time (right). Middle and bottom row: Trace plots for the total number of recombinations (bottom) and for the log-likelihood (middle) for different combinations of *T* and *N* and using 10 million iterations.

#### Example 3.

Typical setting: Comparison across schemes with different sequence lengths and different number of sequences. For all cases *δ* = 236, *θ_s_* = 0.03, *ρ_s_* = 0.005.

Here we compare how the ESS is affected as the length of the sequences becomes larger and as the number of sequences increases. Figure 7 presents the results for lengths of 10Kb, 20Kb and 50Kb. From the plots in the bottom row, the inverse of the ESS appears to be linear as the length of the sequence increases, more specifically, doubling the sequence length halves the ESS. Interestingly, the reduction of ESS for the RJMCMC chain (in blue) when taking into account the running time seems more drastic than the other schemes.

In contrast, increasing the sequence length seems to have a logarithmic effect on the inverse of the ESS as seen in Figure 8; however, when considering running times, the effect becomes linear. For this case, the method with less reduction of ESS is the RJMCMC chain. Thus, RmJMCMC seems to outperform RJMCMC when having larger sequences rather than when having lots of short sequences.

**Figure 7:**
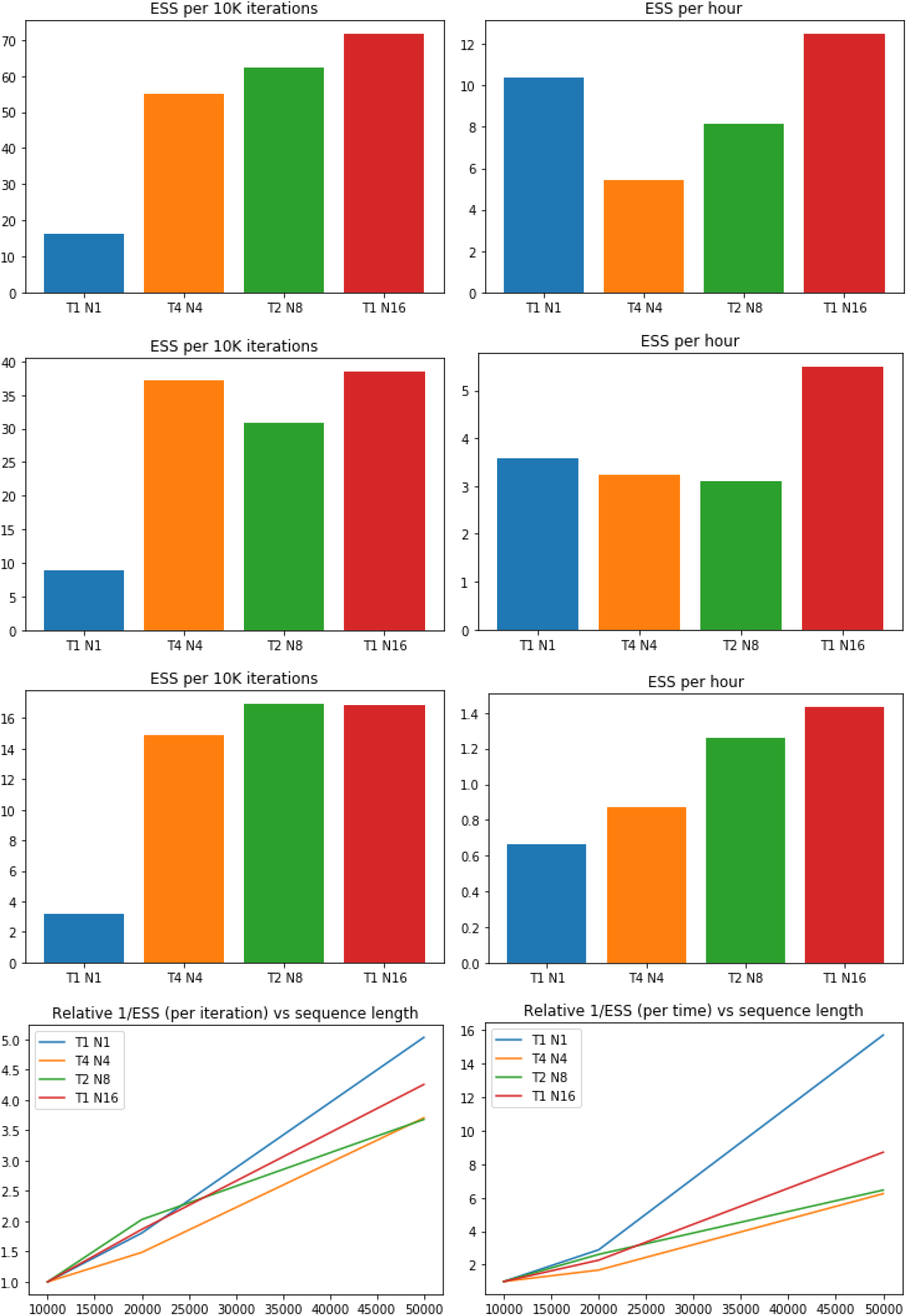
ESS comparison for 10Kb, 20Kb and 50Kb (plots in rows 1-3, respectively) considering 50 sequences in all cases. Bottom plots: increase of the inverse of the ESS as a function of sequence length.

**Figure 8:**
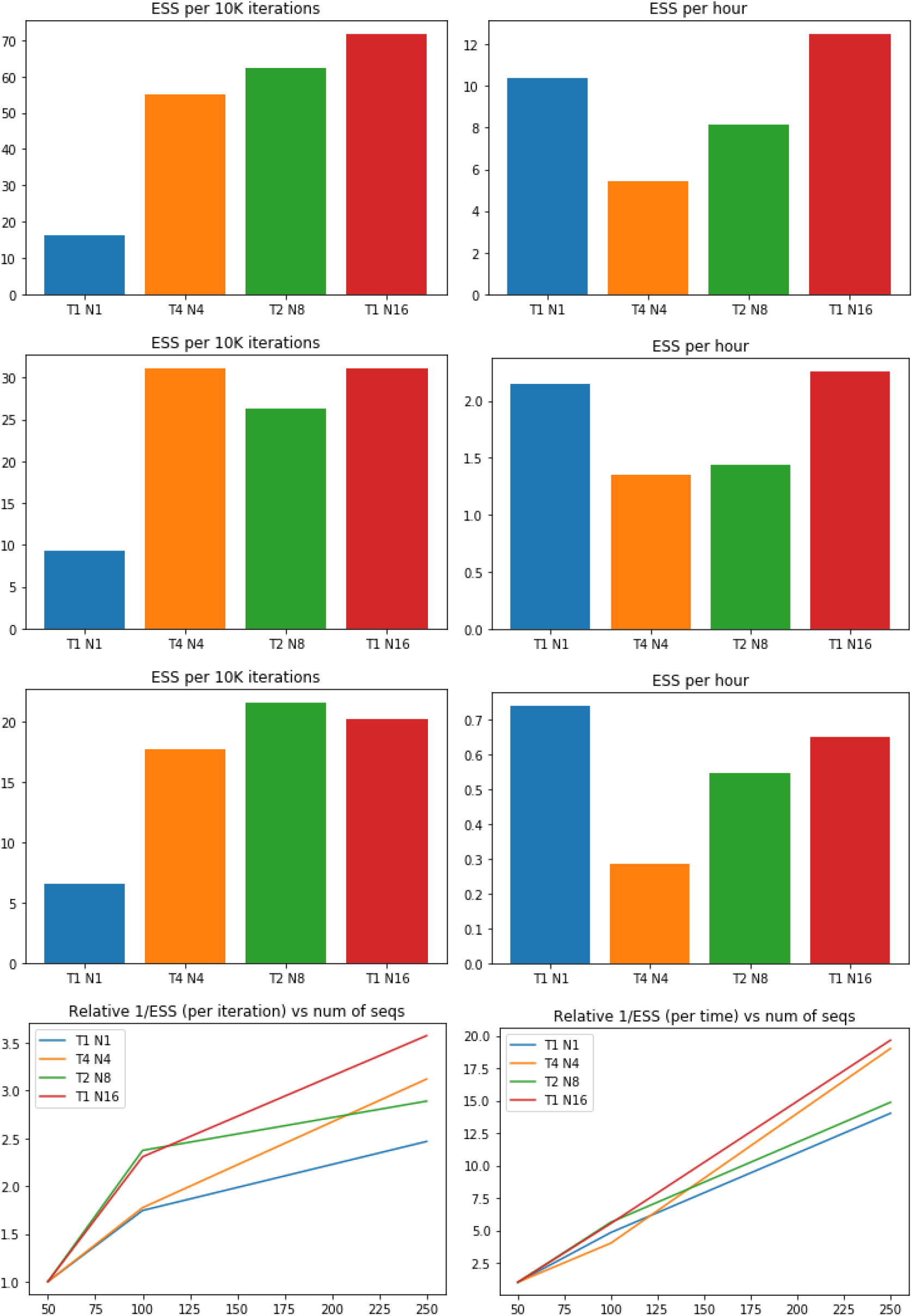
ESS comparison for 50, 100 and 250 sequences (plots in rows 1-3, respectively) considering a length of 10K in all cases. Bottom plots: increase of the inverse of the ESS as a function of sequence length.

### Application to *Escherichia coli* data

Here we present results for a dataset of 27 sequences from *Escherichia coli* on two regions with different recombination intensity; this dataset has been previously studied using ClonalOrigin (Didelot et al., 2012). Similarly as in the simulated examples, we have fixed the clonal genealogy 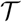 and also some of the global parameters. The specific details on how these fixed values were obtained can be found in Didelot et al. (2012).

#### Example 4.

*E. coli* data on region of length 27.1Kb. The fixed parameters considered here are the clonal genealogy 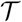 and the mutation rate per site *θ_s_* = 0.0125. Therefore, the inference is performed on the set of recombination events 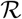 and the global parameters *ρ* and *δ*, i.e. the target distribution arises from (1) by fixing 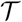 and *θ* accordingly.

Figure 9 presents some preliminary results on a region with a moderate recombination frequency of *ρ_s_* near 0.01. Observe that despite increasing *N* does not seem to improve the value of the ESS adjusted for running time, the log-likelihood trace plot appears to indicate there is some advantage of taking *N* > 1 during the burn-in phase. Interestingly, the trace plots in the bottom row show that when *N* > 1 the chain appears to favour large values for *ρ* during the burn-in stage, whereas for *δ* might take different routes towards the apparent high-posterior region. Clearly, further simulations are needed to rule out whether RmJMCMC offers a clear advantage.

**Figure 9:**
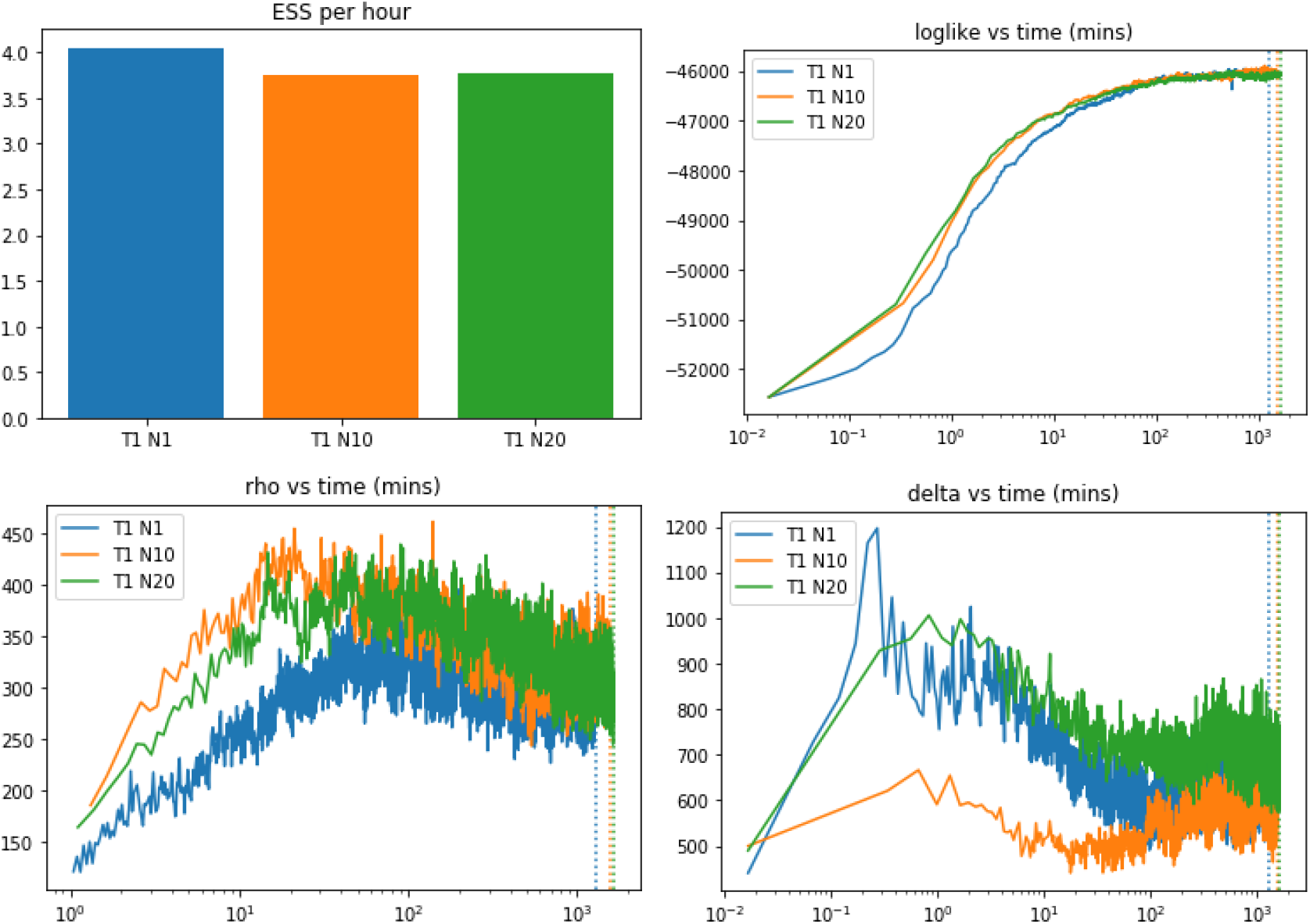
*E. coli* data on region of length 27.1Kb. Top-left: ESS comparison for three different RmJMCMC schemes where *T* = 1; top-right: trace plots for the log-likelihood; bottom-left: trace plots for *ρ*; bottom-right: trace plots for *δ*.

#### Example 5.

*E. coli* data on region of length 3.4Kb. Here we consider a shorter region but for which we know the recombination frequency is higher. The fixed parameters considered once again are the clonal genealogy 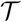 and the mutation rate per site *θ_s_* = 0.0125, we have also fixed the tract length *δ* = 542 due to the high levels of recombination. The inference is thus performed on the set of recombination events 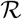 and the global parameter *ρ*, i.e. the target distribution arises from (1) by fixing 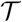, *θ* and *δ* accordingly.

Observe from Figure 10 that this is a more challenging region for the *E. coli* data. In particular, observe that apparently the setting *N* = 12 does not provide an advantage in terms of reaching a high likelihood region in a shorter time; however, when looking at the trace plots for the number of recombination events and for *ρ* it does appear to have an improvement. In this case ESS plots have not been presented since clearly the chain has not reached stationarity, even after nearly 21 days of running the algorithms. Recall that accurate ESS values are based on the fact that the chains have converged. Finally, observe that the recombination rate in this region is much higher than the previous region, with *ρ_s_* reaching values around 0.17. Due to the difficulty of this example, further simulations are needed in order to have a clearer picture of the benefits, if any, from running a RmJMCMC algorithm.

**Figure 10:**
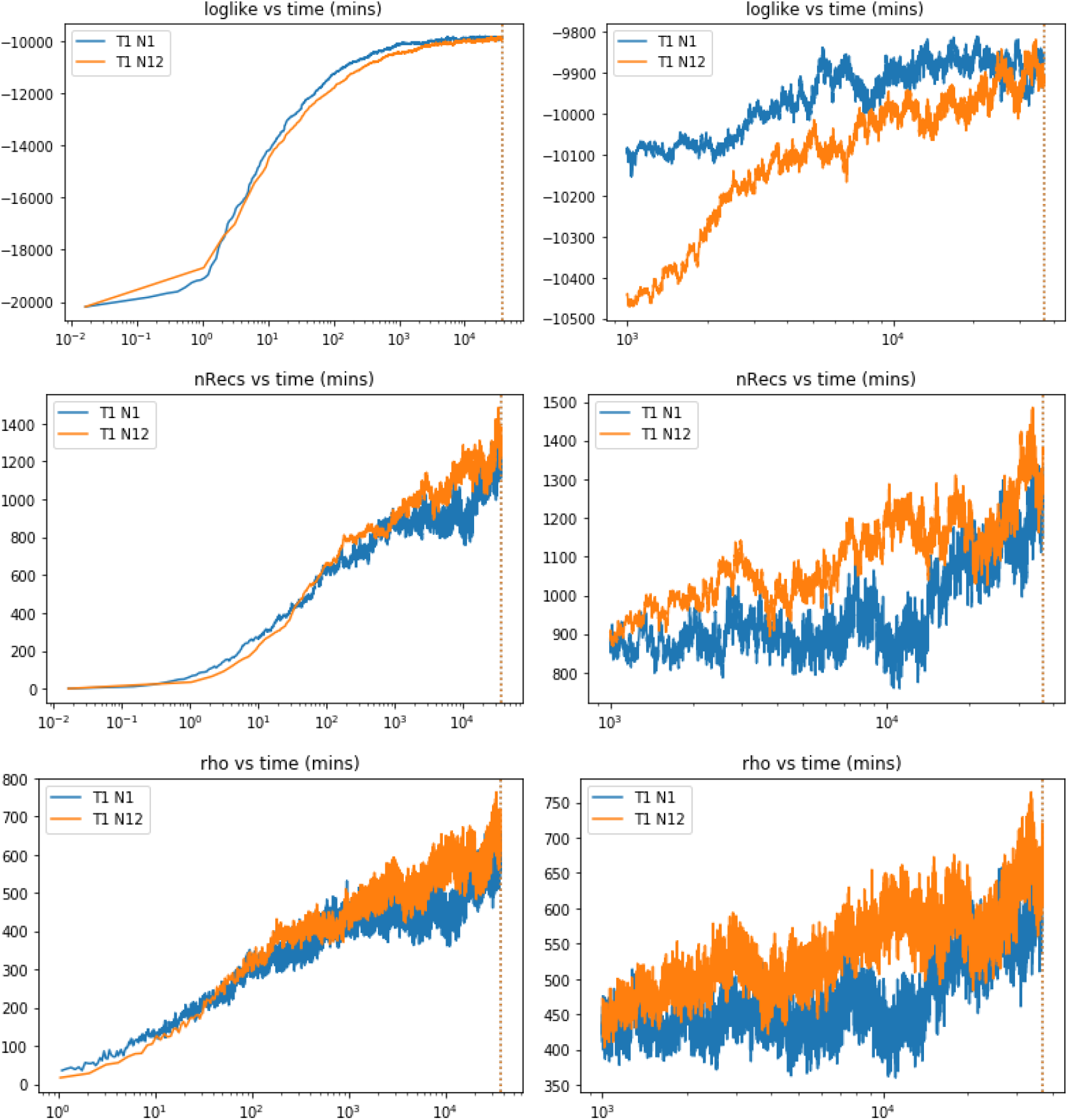
*E. coli* data on region of length 3.4Kb. Top-left: ESS comparison for three different RmJMCMC schemes where *T* = 1; top-right: trace plots for the log-likelihood; bottom-left: trace plots for *ρ*; bottom-right: trace plots for *δ*.

## Discussion

The RmJMCMC algorithm has the potential of speeding up the convergence in the ClonalOrigin model towards a region of high likelihood, hence reducing the number of iterations and running time when compared to a standard RJMCMC. Here the RmJMCMC algorithm has been presented in the context of the ClonalOrigin model, but in principle it could be implemented to other models in population genetics where the dimension of the posterior is not fixed.

The version used in the examples presented here is possibly the simplest one could implement, namely using the prior distribution for proposing and perturbing recombination events. However, other alternatives are possible that may lead to better results, for example splitting or merging existing recombination events, or modifying the prior accordingly if some regions in the tree or in the genome are more likely to recombine. Similarly, different perturbations in the annealing step may be considered since in the presented examples there is no clear advantage when taking *T* > 1. Alternatives include random-walk style proposals for some (or all) of the variables involved in the proposed recombination event (*a_i_, b_i_, x_i_, y_i_*). In this respect, within the annealing process, one should also consider perturbing all of the existing recombination events and not just the recently proposed one.

Finally, it is worth pointing out that the number of cores used in the presented examples was not particularly large; nonetheless we were able to observe significant improvements in some cases. It would be interesting to perform a much larger study where several hundred cores are at hand.

## Supplementary Material

### Details on priors

We use a coalescent tree to represent the clonal genealogy of *n* samples and denote a tree by 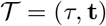, which is composed of a topology *τ* and a vector **t** = (*t*_2_,…, *t_n_*) of branch lengths. A Kingman’s coalescent prior (Kingman, 1982) is assumed for 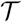, meaning

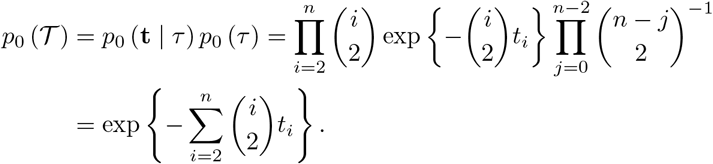

The number of recombination events *R* is assumed, a priori, to be a Poisson random variable with mean 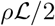, where *ρ* denotes the global recombination rate and 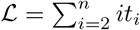 is the total branch length of the tree 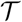. Hence

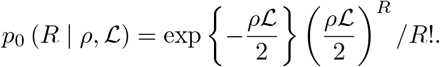

Each recombination event is formed by the following variables: departure and arrival points on the genealogy, and start and end sites on the genome. These four variables, when referring to the *i*-th recombination edge, are denoted by *a_i_, b_i_, x_i_* and *y_i_*, respectively. Both variables *a_i_* and *b_i_* are fully determined by a time and lineage on the tree (denoted respectively by *a_i_*[*t*] and *a_i_*[*l*], respectively), and the chosen priors are those from the construction of an ARG (Wiuf and Hein, 1999). These are a uniform distribution on the tree for *b_i_*, and from this arrival point the edge reconnects at *a_i_* on the tree at a rate proportional to the number of active lineages. We then have

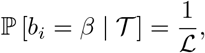

and, assuming that at time *b_i_*[*t*] there are *m* ∈ {2,…, *n*} active lineages and letting 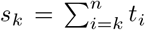, if *u* ∈ (*b_i_*[*t*], *s_m_*)

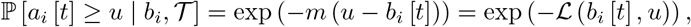

where 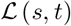 denotes the sum of branch lengths from time *s* to time *t* on the tree. If *u* ∈ (*s_k_*, *s*_*k*−1_) for any *k* ∈ {2,…,*m*}, considering *s*_1_ = ∞,

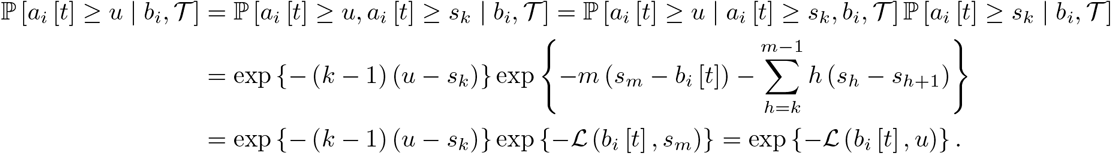

Therefore,

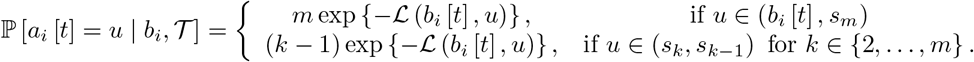

Once *a_i_*[*t*] has been found, the lineage *a_i_*[*t*] on which the recombination event departs is chosen uniformly across the active lineages. Hence

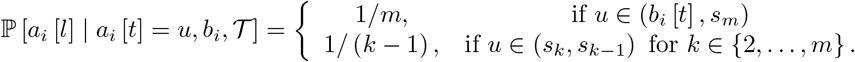

The previous two probabilities lead to

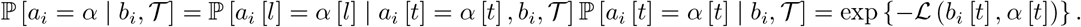

The priors for *x_i_* and *y_i_* are constructed assuming a uniform distribution on the sequence for *x_i_* and a geometric distribution of mean *δ* > 0 for the difference *y_i_* − *x_i_* | *x_i_*. When the sequence is made of *B* blocks comprising a total length of 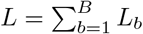, the priors need to be modified accordingly as in (Didelot et al., 2010). Let 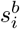 denote the *i*-th site in block the *b*-th block, where *i* ∈ {1,…, *L_b_*} and 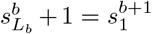, then if *u* is not the first site of a block 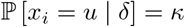 (a constant); whereas if *u* is at the beginning of a block (say 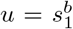) we can use the uncensored variables 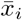 and 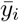 (which denote the true sites where the recombination begins and ends) to derive

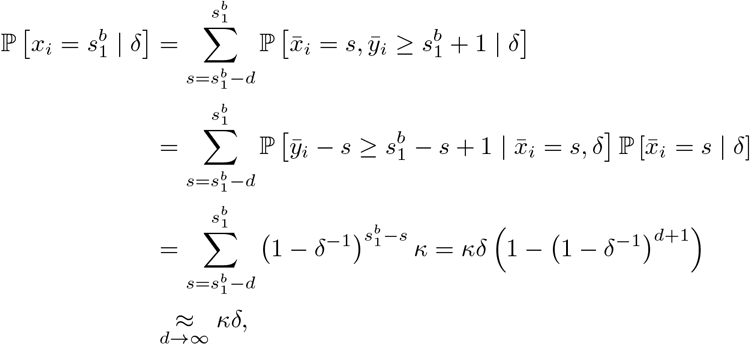

where *d* denotes the number of sites separating the (*b* − 1)-th and *b*-th blocks. Summing over all possible values for *u*, i.e. *B* sites at the beginning of blocks and *L* − *B* in other regions, we get (*L* − *B*) *κ* + *Bκδ* = 1; this implies *κ* = 1/ (*δB* + *L* − *B*), leading to

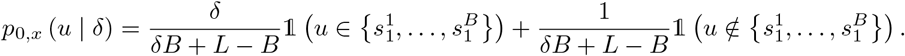

In order to find the density of *y_i_* we use the fact that 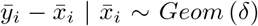 on the positive integers. We then have for *v* ≥ *u* and 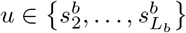

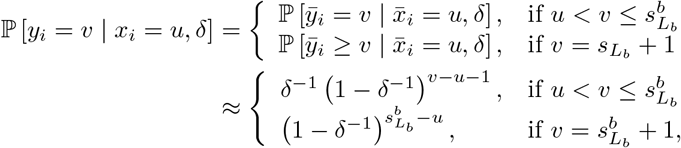

and similarly if 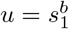

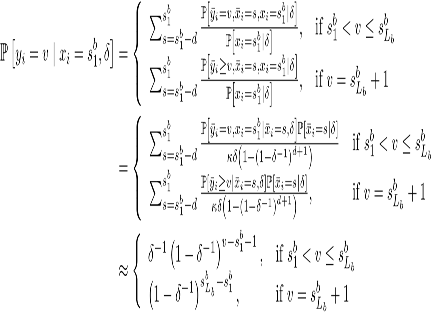

Hence,

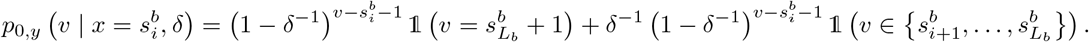

The previous equations fully define the prior to be used for the set of recombination events, denoted by 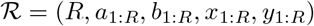, i.e.

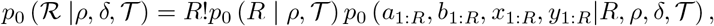

where the factorial comes from ordering the recombination events (e.g. ordering according to one of the continuous variables **x** or **y**). Assuming recombination events occur independently given the genealogy 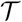 and introducing priors for *ρ* and *δ*, we obtain

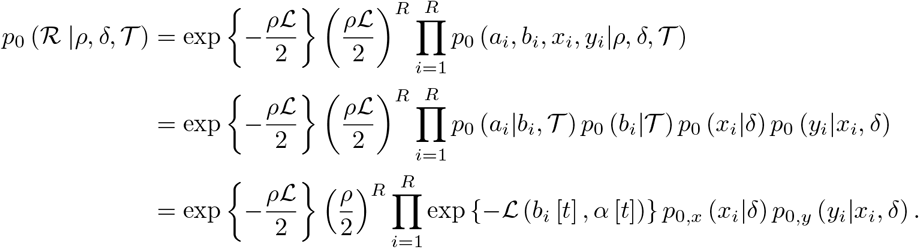

### Likelihood computation

Recall that the vector 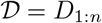 corresponds to the *n* observed DNA sequences each composed of *L* sites, we denote the *i*-th site in the *k*-th sequence by 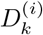. For computing the likelihood function at the *i*-th site, the local tree 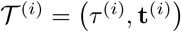 is constructed as a function of the clonal genealogy 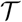 and the set of recombination events 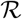. From this local tree with *n* leaves, let 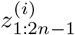 denote the whole set of ordered nodes where of 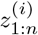 denotes the leaves of the tree. For *k* ≥ *n* +1, let 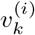 and 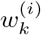 be the left and right children nodes, respectively, of the node 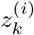. The likelihood of the mutation rate *θ* and the local tree 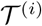 is given by the recursive formula

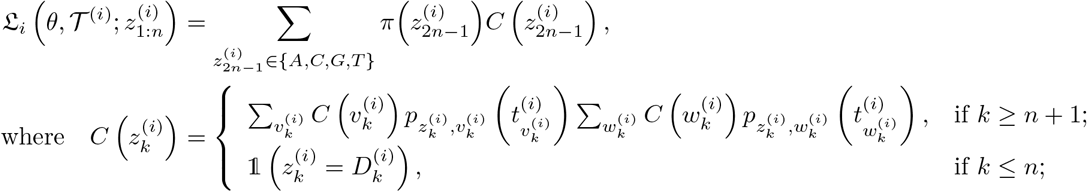

and *p_z,w_*(*t*) denotes the transition probability from the basis *z* to the basis *w* in *t* units of time, and *π* corresponds to the limiting probability of such transitions.

For the JC69 model (Jukes and Cantor, 1969), we have *π*(*z*) = 0.25 for each *z* ∈ {*A, C, G, T*} and

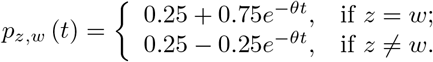

Hence, assuming independence across the *L* different sites in all the sequences we obtain

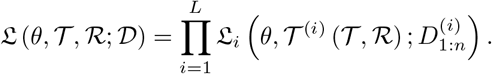

### RJMCMC as an importance estimator within MCMC

To keep notation simple we remove the variables 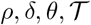 and the data 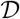 from the conditional distributions, i.e. we write 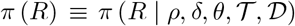. In order to jump up in dimension, i.e. to go from *R* recombination events to *R* + 1, we must create four variables *a_R_, b_R_, x_R_, y_R_*. This is done using an auxiliary distribution, for example the prior for such variables, which we denote by 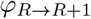 and that in principle could also depend on the data or other parameters. Similarly, in order to perform a jump down in dimension we need a way to merge or remove recombination events, this is done using the distribution 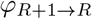. Letting 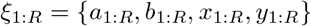, the auxiliary *φ′s* allow us to transform *ξ*_1:*R*_ into 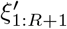 and vice-versa, where *ξ*_1:*R*_ is not necessarily the same as 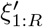. For simplicity, suppose that the resulting one-to-one transformation (which we denote by *T*_*R*→*R*+1_) has a Jacobian with determinant equal to 1, then

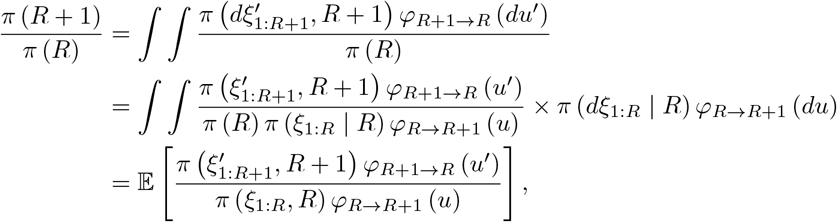

the above expectation is taken on the variables 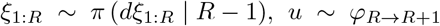, and where 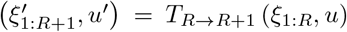. The ratio inside the previous expectation is exactly the ratio used in the RJMCMC algorithm, which serves as an estimate for the ratio of the “ideal” unfeasible scenario in which we could explore the posterior of *R* without the need of *ξ*_1:*R*_.

### Annealing Moves

The full algorithm requires the introduction of the upwards and downwards annealed moves. For this, we require the introduction of auxiliary distributions, the first 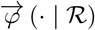 is for generating a new recombination event, i.e. for moving upwards; whereas the second 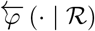 is for selecting an index when deleting a recombination event. Moreover, these two distributions might also depend on 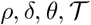 and/or 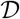. Simple choices are the prior of a single recombination event (*a,b,x,y*) for 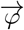 (described above), and a uniform distribution on {1,…, *R*} for 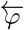 assuming there are *R* active recombination events. We now present the upwards and downwards moves.

Despite the previous algorithms being very similar, they have important differences. For instance, the upwards move requires as input a newly created recombination event, whereas for the downwards this requirement is replaced with the need for an index. Also, even though the sequence 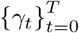 is required in both cases, the construction of *η_t_* is different depending on the type of move. We want to stress that this construction is essential for obtaining a valid algorithm, otherwise the procedure could target something different to the desired posterior. A simple choice for the sequence if *γ_t_* = *t*/*T*, which implies *γ*_*T*−*t*_ = 1 − *γ_t_*.

Finally, we emphasise that in both algorithms, the variables 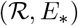 can be obtained from 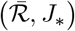 and viceversa; this means that ratios of the form *η*_*t*+1_/*η_t_* can be solely expressed in terms of 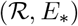 or 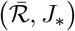. Similarly, when perturbing 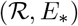 using a MCMC process we are implicitly perturbing the corresponding 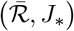. In addition to this, the outputs are not that different since one set of variables can be obtained from the other. In both algorithms we chose to work with 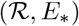 and only compute the corresponding pair 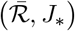 when needed.

#### Algorithm 2 Upwards Annealed Move

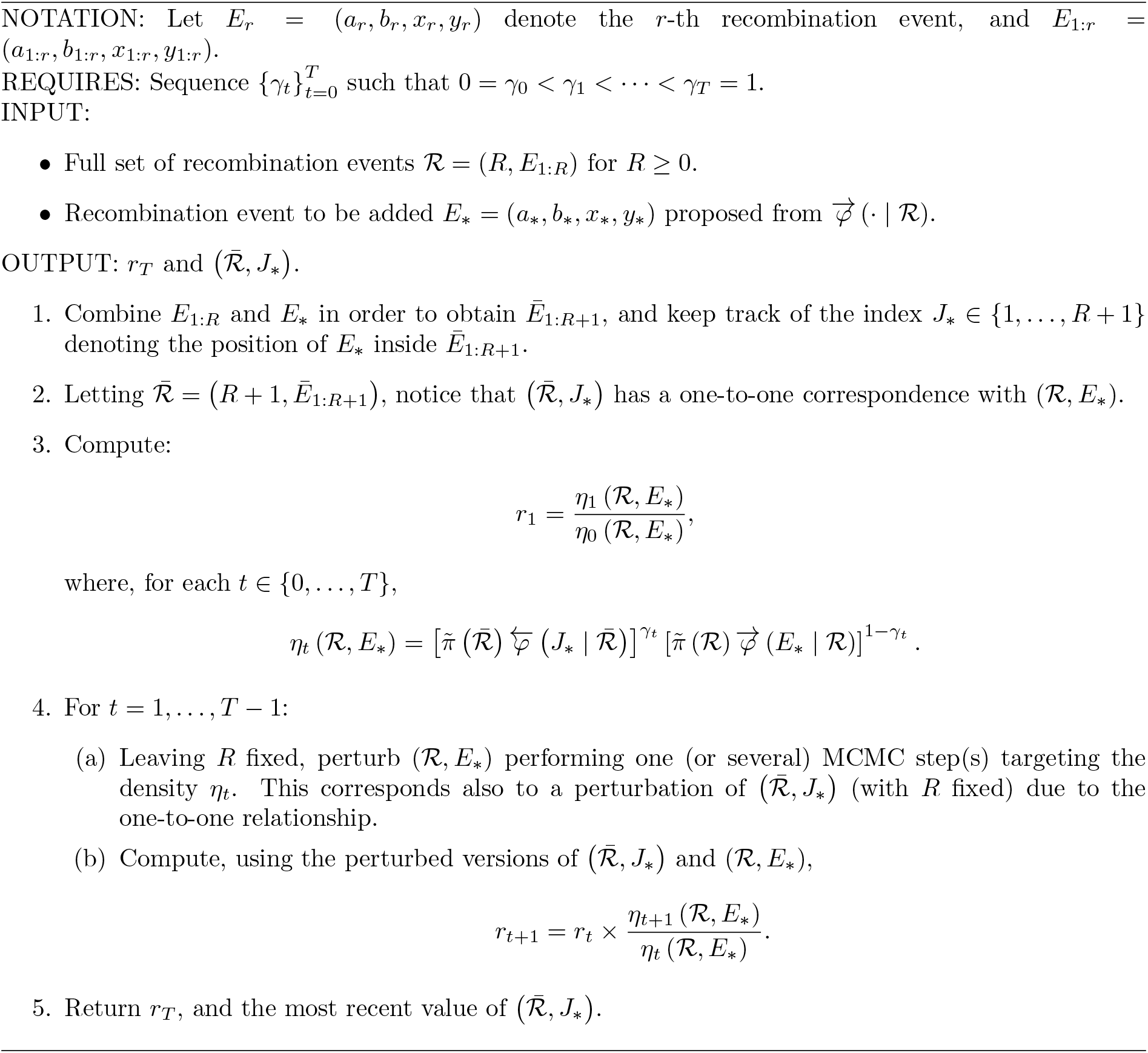

#### Algorithm 3 Downwards Annealed Move

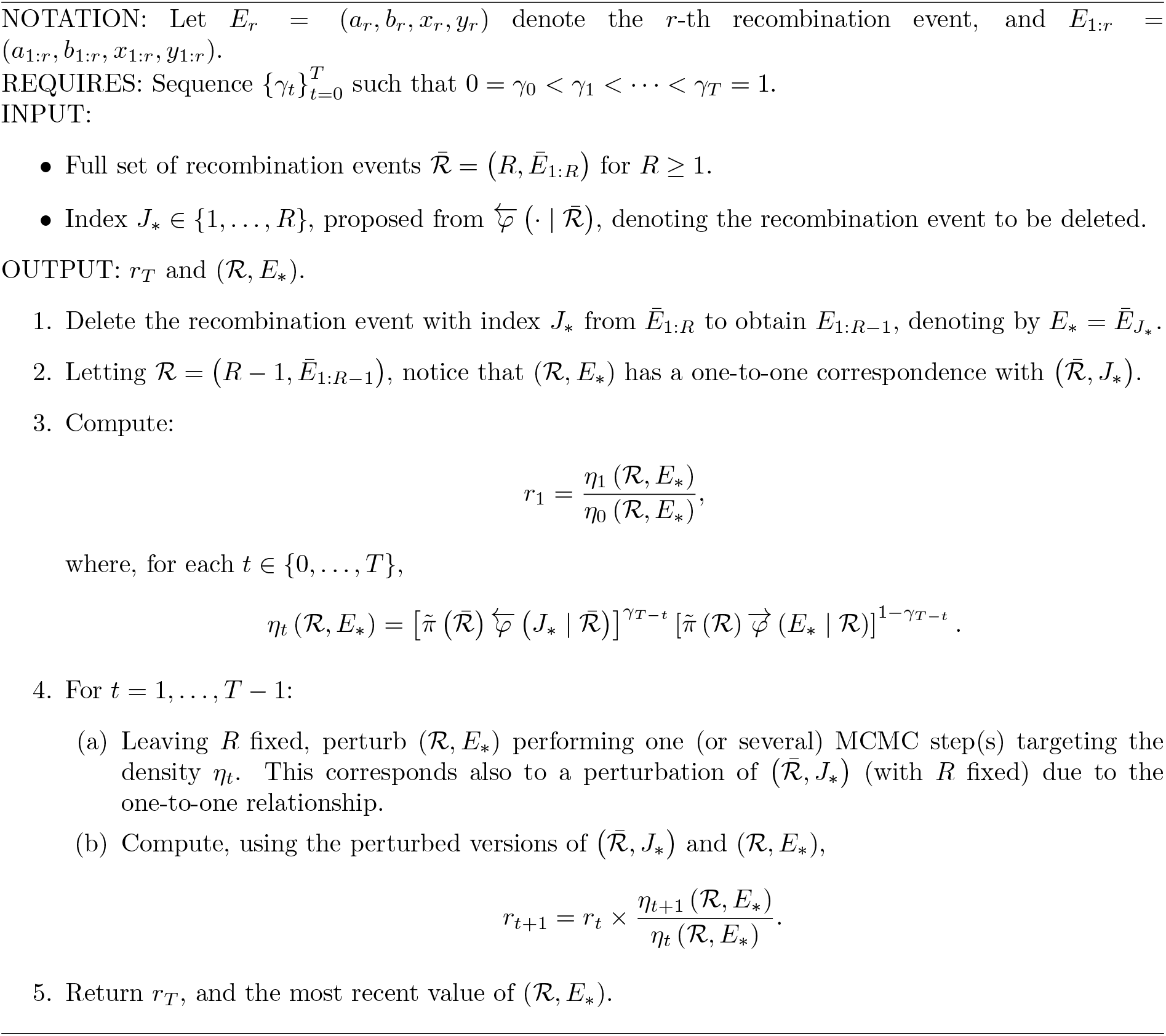

